# Inflammation induced T_h_17 cells synergize with the inflammation-trained microbiota to mediate host-resiliency against intestinal injury

**DOI:** 10.1101/2024.03.25.586435

**Authors:** JL Golob, G Hou, BJ Swanson, S Bishu, H Grasberger, M El Zataari, A Lee, J Kao, N Kamada, S Bishu

## Abstract

**Background and Aims:** Inflammation can generate pathogenic T_h_17 cells and cause a inflammatory dysbiosis. In the context of Inflammatory Bowel Disease (IBD) these inflammatory T_h_17 cells and dysbiotic microbiota may perpetuate injury to intestinal epithelial cells (IECs). However, many models of IBD like T-cell transfer colitis and IL-10^-/-^ mice rely on the absence of regulatory pathways, so it is difficult to tell if inflammation can also induce protective T_h_17 cells.

**Methods:** We subjected C57BL6, RAG1^-/-^ or J_H_^-/-^ mice to systemic or gastrointestinal (GI) *Citrobacter rodentium* (*Cr*). Mice were then subject to 2.5% dextran sodium sulfate to cause epithelial injury. Fecal microbiota transfer was performed by bedding transfer and co-housing. Flow cytometry, qPCR, 16s sequencing and histology were used to assess parameters.

**Results:** Transient inflammation with GI but not systemic *Cr* was protective from subsequent intestinal injury. This was replicated with sequential DSS collectively indicating that transient inflammation provides tissue-specific protection. Inflammatory T_h_17 cells that have a tissue resident memory signature expanded in the intestine. Experiments with reconstituted RAG1^-/-^, J_H_^-/-^ mice and cell trafficking inhibitors showed that inflammation induced T_h_17 cells were required for protection. Fecal microbiota transfer showed that the inflammation-trained microbiota was necessary for protection, likely by maintaining protective T_h_17 cells *in situ*.

**Conclusion:** Inflammation can generate protective T_h_17 cells which synergize with the inflammation-trained microbiota to provide host resiliency against subsequent injury, indicating that inflammation induced T_h_17 tissue resident memory T cells are heterogenous and contain protective subsets.

## INTRODUCTION

Inflammatory Bowel Disease (IBD), encompassing Crohn’s Disease and Ulcerative Colitis are incurable conditions with high morbidity and economic burden. While the exact cause of IBD is unknown, IBD is thought to be due to dysregulated immune responses to dysbiotic gut microbiota ^1–3^. On a tissue level, intestinal epithelial cell (IEC) injury is thought to lead to the loss of tolerance to self and commensals and perpetuate dysbiosis, begetting further injury.

T-cells, which are a major part of host immunity, are strongly linked with IBD. Genome-wide association studies (GWAS) in IBD show that many risk conferring single-nucleotide polymorphisms (SNPs) are either upstream of or are directly expressed by CD4^+^ T-cells ^4–8^. Within CD4^+^ T-cells, the IL-17A producing subset (‘T_h_17’ cells), are even more strongly implicated in the pathogenesis of IBD. GWAS and candidate gene studies have identified SNPs in multiple T_h_17 pathway genes including *CARD9*, *IL23R*, *IL21*, *RORC*, *IL12B*, and *JAK2* ^4^. Though many of these pathways have pleiotropic functions *IL23R* and *RORC* are specifically linked with T_h_17 cells ^5, 9–11^. Adding to human studies, murine models of colitis including IL-10^-/-^ and T-cell transfer strongly link the IL-23:T_h_17 pathway with IBD ^7, 12–14^. Evidencing the strength of this association, biologics like Ustekinumab, Risankizumab and Guselkumab which target IL-23:T_h_17 pathways are efficacious in IBD ^15–17^.

Adding another layer of complexity, terminally differentiated CD4^+^ T-cells have highly specialized transcriptional signatures and phenotypes specific to the anatomic compartment(s) they reside in ^18, 19^. CD4^+^ T effector-memory (T_EM_) primarily reside in the systemic circulation, though they can migrate to peripheral sites when activated ^19^. In tissues at the host-microbiome interface like the skin, genitourinary, pulmonary, and gastrointestinal (G) tracts, so-called tissue resident memory (T_RM_) T-cells are the major T-cell subset ^2, 18–26^. T_RM_ can be CD8^+^ or CD4^+^, of which much less is known about CD4^+^ T_RM_ ^23–25, 27, 28^. Once formed, T_RM_ are largely tissue restricted with little systemic re-circulation, which is ensured by downregulation of cell trafficking receptors like sphingosine-1-phosphate receptor 1 (S1PR1), CD62L and CCR7 that would allow for tissue egress ^27, 29, 30^. T_RM_ function as a first line of adaptive defense against invading pathogens *in situ*^18, 20, 21^. Beyond host defense, we and others have linked CD4^+^ T_RM_ with human IBD and CD4^+^ T_RM_ can causally induce colitis in murine models ^26, 31,32^. Amid these points, it is important to note that not all CD4^+^ T-cells are pathogenic. Some can be protective, including anti-inflammatory subsets of T_h_17 cells and the canonically protective T-regulatory cells^33–35^. Whether some CD4^+^ T_RM_ can be protective is unknown.

In exploring these questions, we unexpectedly found that transient GI inflammation is protective against subsequent IEC injury. Systemic insults did not confer IEC protection, indicating that protection is encoded in the intestinal environment. Using IL-17 GFP fate-mapping (IL-17^FM^) mice and multiple approaches including RAG1^-/-^ and J_H_^-/-^ mice and pharmacologic inhibitors of cell trafficking, we found that inflammation induced intestinal T_h_17 cells exhibit a T_RM_ signature and are required for protection. Furthermore, consistent with tissue-level encoding, transient inflammation durably altered the gut microbiota. IEC protection could be ameliorated by disrupting this inflammation-trained microbiota, but protection could not be transferred by fecal microbiota transplantation (FMT). Therefore, our data collectively show that transient inflammation is protective from subsequent IEC injury via a pathway coupling inflammation induced T_h_17 T_RM_ with the inflammation-trained microbiota.

## RESULTS

### Transient intestinal inflammation can protect from subsequent epithelial injury

As an initial model of transient GI inflammation we used the bacteria *Citrobacter rodentium* (*Cr*), which causes a self-limiting infection with epithelial damage ^36^. We choose *Cr* because the immune responses to *Cr* are well characterized, and because *Cr* is completely cleared in all immune-competent mice, leaving them in a basal state without any impairments ^12, 37–40^. We took advantage of the fact that *Cr* strongly induces T_h_17 and used IL-17A GFP fate-mapping (IL-17^FM^) mice to track T_h_17 indefinitely ^24^. Early (day < 5) immunity to *Cr* depends on innate lymphoid cells (ILCs) and late immunity depends on CD4^+^ T-cells and humoral immunity. Other T-cell subsets like CD8^+^, NKT, and γδ T-cells are dispensable for immunity to *Cr* ^12, 39, 41^.

We subjected IL-17^FM^ mice to gastrointestinal (GI) *Cr* or PBS via oral gavage (**Fig 1A**). GI *Cr* generates intestinal and systemic immune responses. To ascertain intestine-specific vs systemic responses, we included intravenous (IV) *Cr* because it only generates systemic responses ^39, 42^. Mice cleared mucosal and luminal *Cr* in 4-5 weeks in our facilities and were left unmanipulated for at least 2 more weeks, at which time they are healthy and indistinguishable from uninfected controls ^25^ (**Sup Fig 1A**). Six weeks post-infection (p.i.) (which is 2 weeks after recovery from *Cr*), mice were administered 2.5% dextran sodium sulfate (DSS) to cause intestinal epithelial cell (IEC) injury (**Fig 1A**). *Cr*-induced T_h_17, which are considered pro-inflammatory, become abundant in the intestine by this time ^23–25^. So, we expected that transient inflammation would exacerbate subsequent IEC injury because of the expansion of inflammatory T_h_17 cells in the intestine. Surprisingly however, transient inflammation was protective against subsequent IEC injury. Mice that had recovered from transient gut inflammation with *Cr* had less weight loss, milder disease and exhibited significantly longer survival that mice that had recovered from IV *Cr* or PBS controls (**Figs 1B-D, Sup Fig 1B**). The protection was associated with less epithelial damage and fewer immune cell infiltrates (**Fig 1E**). Although transient inflammation with GI *Cr* is protective, these mice still up regulate pro-inflammatory T_h_17 genes with DSS, suggesting that inflammatory T_h_17 cells are activated during IEC injury (**Sup Fig 1C**). Importantly, only GI Cr is protective, and IV Cr is not (**Fig 1**). GI *Cr* causes intestinal inflammation, but IV *Cr* does not, meaning that the anatomic site of inflammation and protection are the same and protection after inflammation is a tissue-specific response.

**Figure 1.**
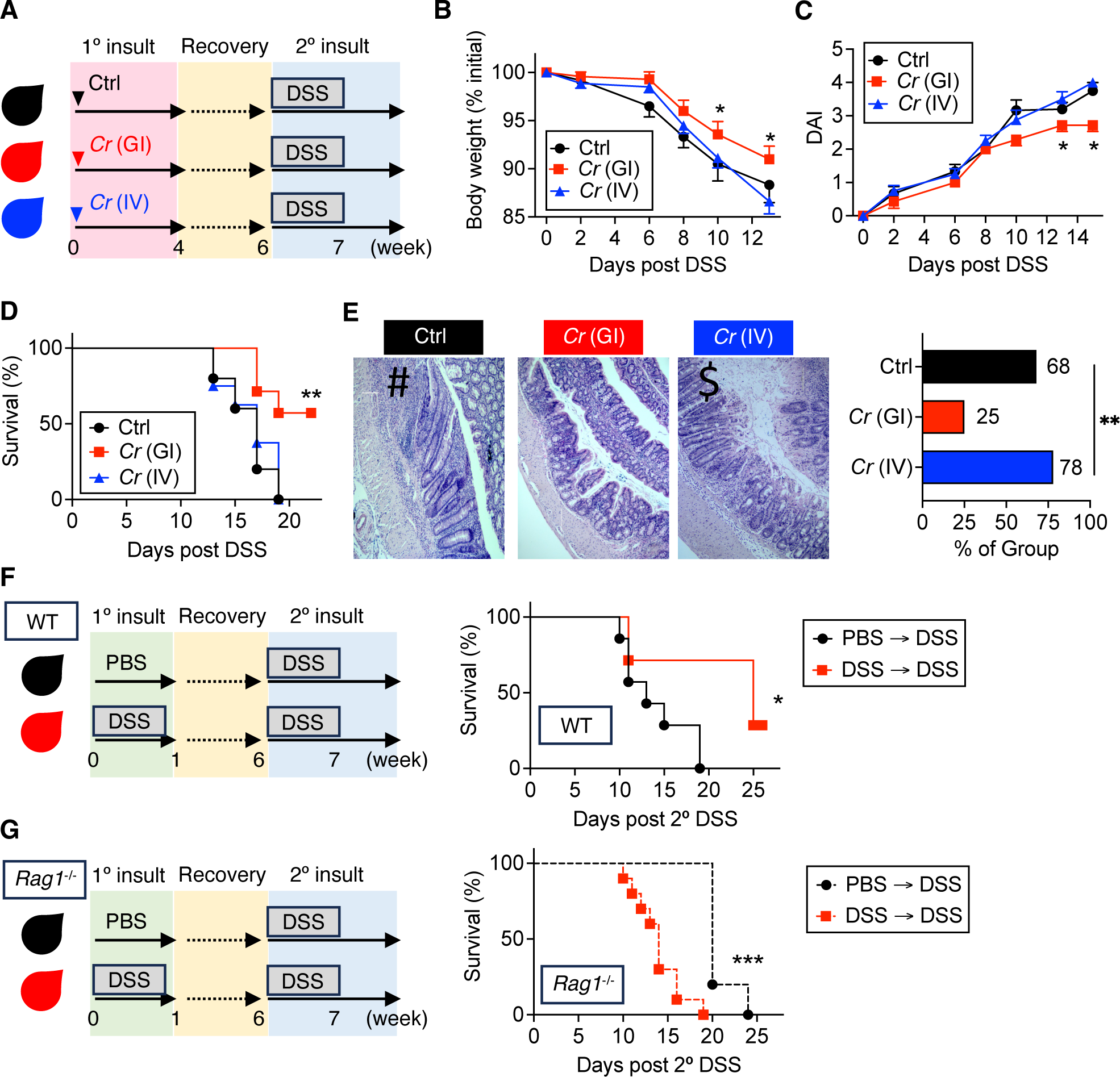
Transient intestinal inflammation can protect from subsequent epithelial injury. **A)** Mice were subject to *C. rodentium* (*Cr*) infection via oral gavage (gastrointestinal [GI]) or intravenous (IV) inoculation and allowed to recover, which takes 4-5 weeks. Six weeks post infection, mice were subject to 2.5% DSS and **(B)** weight change, **(C)** disease activity index (DAI), and **(D)** survival were measured. **E)** Histology showing inflammatory infiltrates (#) and ulcers ($) (left panel) and inflammation as a composite of immune cell infiltration and epithelial injury (right panel) were scored. **F)** Wild-type or **(G)** *Rag*1^-/-^ mice were subject to PBS or 2.5% DSS for 7 days, then allowed to recover. At week 6, all mice were subject to 2.5% DSS and survival was measured (right panels). * to *** p < 0.01-0.0001 by ANOVA (B, C, E) or Kaplan-Meier (D, F, G). Data are representative of 6 experimental replicates, with 3-5 mice per group, per experiment.

As a second system of transient inflammation, we subjected mice to DSS for 7 days and allowed them to recover. Six weeks later, we subjected them to a second cycle of DSS (**Fig 1F**, top panel). Mice recover from DSS in 2 weeks, so this time is ample to return to baseline ^43^. Like transient inflammation from GI *Cr*, one cycle of DSS is protective against a second cycle (**Fig 1F**, bottom panel**, Sup Fig 1D**) which is a familiar result to those who use chronic DSS models.

The 6-week lag between the initial inflammation and subsequent injury suggests a memory response, wherein inflammation primes the organism to be resilient against subsequent injury. For *Cr* this is long-enough that ILCs are unlikely to be involved ^40, 41^. We therefore focused on adaptive immunity which is strongly induced after gut inflammation. To test the role of adaptive immunity, we used RAG1^-/-^ mice which lack B- and T-cells. Furthermore, since RAG1^-/-^ mice succumb to *Cr*, we used the sequential DSS model (**Fig 1G**). In contrast to wild-type (WT) mice, transient inflammation did not protect RAG1^-/-^ mice from IEC injury, indicating that adaptive immunity is required (**Fig 1G**). These data show that transient GI inflammation can protect from subsequent IEC injury and that protection is a tissue-specific response. These data therefore imply that protection requires tissue-specific adaptive immunity.

### Transient inflammation generates tissue-specific T_h_17 subtypes

Our data indicates that tissue-specific memory responses are required for IEC protection after transient inflammation (**Fig 1)**. Intestinal adaptive immunity clearly fulfils these criteria and conceptually fits our data. We therefore explored this possibility.

To examine how intestinal immune populations change after transient inflammation, we subjected IL-17^FM^ mice to GI or IV *Cr* or (oral) PBS and analyzed intestinal lamina propria mononuclear cells (LPMCs) at week 6 p.i. with flow cytometry (FCS) (**Fig 2A**). To broadly examine intestinal immune populations we used a simple strategy based on the expression of CD3e, CD4 and CD8b. This scheme covers most LPMC subsets at the cost of granular detail. For example, ILCs, B-cells and most antigen presenting cells are in the CD3e^-^ fraction and the CD3e^+^CD4^-^CD8b^-^ fractions contain all non-T_h_ and non-T_c_ subsets like NKT, MAIT and gd T-cells. PBS gavaged controls had a preponderance of CD3e^+^CD4^-^CD8b^-^ fractions in the intestine. This distribution shifted strongly towards CD4^+^ T-cells specifically with GI *Cr* (**Fig 2B, C**). We therefore focused on CD4^+^ T-cells.

**Figure 2.**
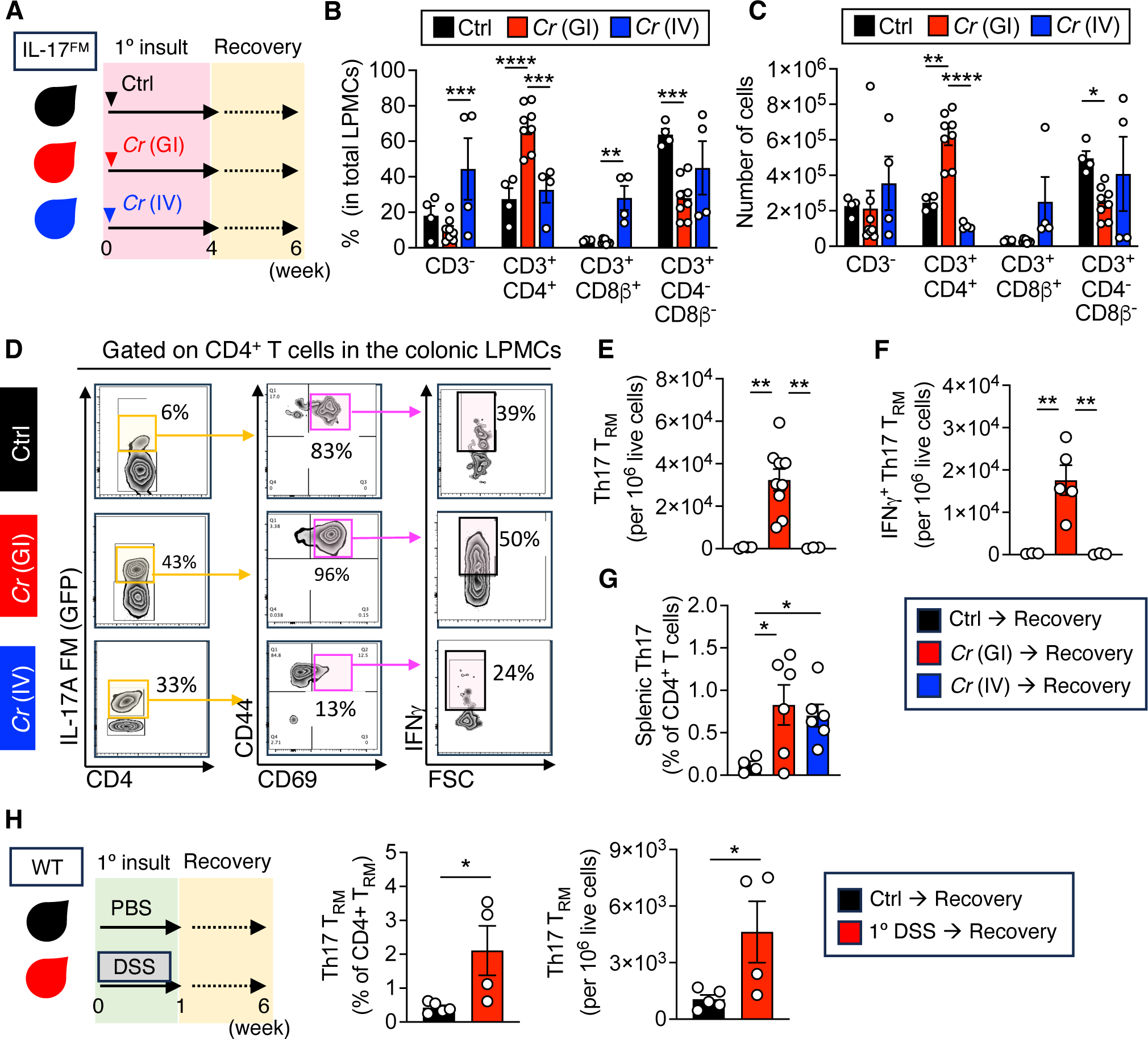
Transient inflammation generates tissue-specific T_h_17 subtypes. **A)** IL-17^FM^ mice, which permanently activate GFP upon IL-17 expression, were subjected to GI or iv *Cr* or PBS. They were allowed to recover and sacrificed at week 6. Live lamina propria mononuclear cells (LPMCs) were gated from mice in (A) into groups as indicated and their **(B)** relative fractions and **(C)** absolute numbers were quantified. **D)** Intestinal CD4^+^ T-cells from mice in (A) were gated as shown, with further gating on GFP^+^ (T_h_17) (middle and right columns). **E)** T_h_17 T_RM_ defined as CD4^+^GFP^+^CD44^+^CD69^+^ from (D) and **F)** IFNɣ^+^ T_h_17 T_RM_ were enumerated. **G)** The fractions of T_h_17 T-effector memory cells (CD4^+^GFP^+^CD44^+^CD69^-^) in spleen of mice in (A) were quantified. **H)** Wild-type mice were subject to PBS or 2.5% DSS for 1 week and then allowed to recover. LPMCs were assessed by flow cytometry at week 6 and the fraction (left panel) and absolute numbers (right panel) of T_h_17 T_RM_ was quantified * to **** p < 0.01-0.00001 by ANOVA (B-G) or students t-test (H). Data are representative of 2-6 experimental replicates, with 3-9 mice per group, per experiment.

It is important to clarify nomenclature at this point. Because this is an IL-17^FM^ system, GFP^+^CD4^+^ T-cell fractions encompass CD4^+^ T-cells that (1) actively produce IL-17A (‘current’ T_h_17 cells) or (2) those that previously made, but are not currently making, IL-17A (‘*ex*-T_h_17’ cells) ^10, 44^. There is no consensus on the nomenclature of these cells nor a clear definition of what it means to be ‘*ex*-‘, but a recent single-cell sequencing study of *ex*- and current T_h_17 cells found that they have a nearly indistinguishable transcriptional profile ^10^. This implies that T_h_17 cells retain their ‘T_h_17-ness’ even if they are not actively producing IL-17A. So, for simplicity we will consider GFP^+^CD4^+^ T-cells as T_h_17 cells unless otherwise noted.

Both IV and GI *Cr* generate GFP^+^CD4^+^ T-cells. However, IV *Cr* generates T_h_17 T-effector memory (T_EM_) cells (GFP^+^CD4^+^CD44^+^CD69^-^) which predominately stay in the circulation. In contrast, GI *Cr* generates T_h_17 cells that exhibit a T_RM_ profile characterized by the expression of CD44 and CD69 (**Figs 2D, E**). Substantial fractions of these T_h_17 cells also produce interferon (IFN)g (**Fig 2F**). Thus GI but not IV *Cr* generates T_h_17 cells that are intestine restricted (i.e. T_RM_) which is consistent with our prior work and that of others ^23–25^. T_h_17 expand in the spleen with both IV and GI *Cr*, verifying that both routes of inoculation generate systemic responses (**Fig 2G**). Consistent with these data, one cycle of DSS also generates T_h_17 cells with a T_RM_ signature in the intestine, though to a lesser degree than with GI *Cr* (**Fig 2H**).

Finally, to determine the fraction of intestinal T_h_17 which are actively producing IL-17A, we used IL-17A conjugated to APC so that IL-17A producing cells will be GFP^+^APC^+^. In mice that had cleared GI *Cr*, about 15% of intestinal T_h_17 are actively producing IL-17A. This data also verifies the fidelity of the IL-17^FM^ mice since < 4% of GFP^-^ fractions are IL-17APC^+^, which is probably the basal autofluorescence rate (since this level is similar between GI *Cr* and PBS) (**Sup Fig 2A, B**). These data show that transient inflammation leads to tissue-specific T_h_17 subtypes, and that IEC protection strongly correlates with the expansion of intestinal T_h_17 cells that exhibit a T_RM_ signature. This result is surprising because T_h_17 cells are thought to be primarily pathogenic. However, it is possible that a subset of GI *Cr* induced T_h_17 cells are protective, since even T_h_17 cells have non-pathogenic subtypes ^5, 35^.

### Inflammation induced T_h_17 cells are required for protection after transient inflammation

Our data finds that IEC protection is associated with expansion of inflammation induced T_h_17 cells in the intestine. These T_h_17 cells exhibit a tissue-restricted (T_RM_) signature which is consistent with the finding that protection is a tissue-specific phenomena (**Fig 1**). We thus considered the possibility that the inflammation induced T_h_17 may mediate protection.

The inflammation induced T_h_17 cells exhibit a T_RM_ signature. Testing the role of T_RM_ is experimentally challenging because T_RM_ are disconnected from the circulation, so once formed, they cannot be deleted with antibodies or methods like CD4-cre/ERT2. Cell transfer is also ineffective because colonic T_RM_ downregulate receptors that allow for their transit from blood to tissue, like sphingosine 1 phosphate receptor 1 (S1PR1) and CCR7 ^20, 27, 28, 30^. Thus, transferred colonic T_RM_ never home to the colon in recipient mice. These features of T_RM_, which we verified in our model, make them challenging to study. Therefore, to test whether inflammation induced T_h_17 could be protective, we used three complimentary approaches, namely J_H_^-/-^ mice, reconstitution of RAG1^-/-^ mice, and the agent FTY 720.

J_H_^-/-^ mice lack mature B-cells and humoral immunity but have an intact T-cell compartment ^45^. We reasoned that if *Cr* induced T_h_17 cells are required, then J_H_^-/-^ mice would be protected from IEC injury. However, J_H_^-/-^ mice succumb to GI *Cr* because humoral immunity is required to clear *Cr* ^39, 46^. J_H_^-/-^ mice can be rescued by administering convalescent serum from wild-type (WT) mice that have cleared *Cr*, which contains anti-*Cr* IgG ^39^.

We subjected J_H_^-/-^ mice to GI *Cr* or (GI) PBS concurrent with convalescent serum (**Fig 3A**). To ensure that J_H_^-/-^ mice that received convalescent serum cleared *Cr*, we enumerated *Cr* colony forming units from feces (**Sup Fig 3A, B**). We also verified that circulating convalescent serum was lost from J_H_^-/-^ mice at the time of administering DSS, to ensure we were only testing the role of *Cr*-induced T_h_17 (**Sup Fig 3C**). Thus, these mice have inflammation induced T_h_17 cells (which are predominately T_RM_), but lack inflammation induced humoral immunity (**Fig 3A**). Consistent with the possibility that inflammation induced T_h_17 mediate protection, J_H_^-/-^ mice that had recovered from transient inflammation were protected and had longer survival relative to PBS counterparts (**Fig 3B-D**).

**Figure 3.**
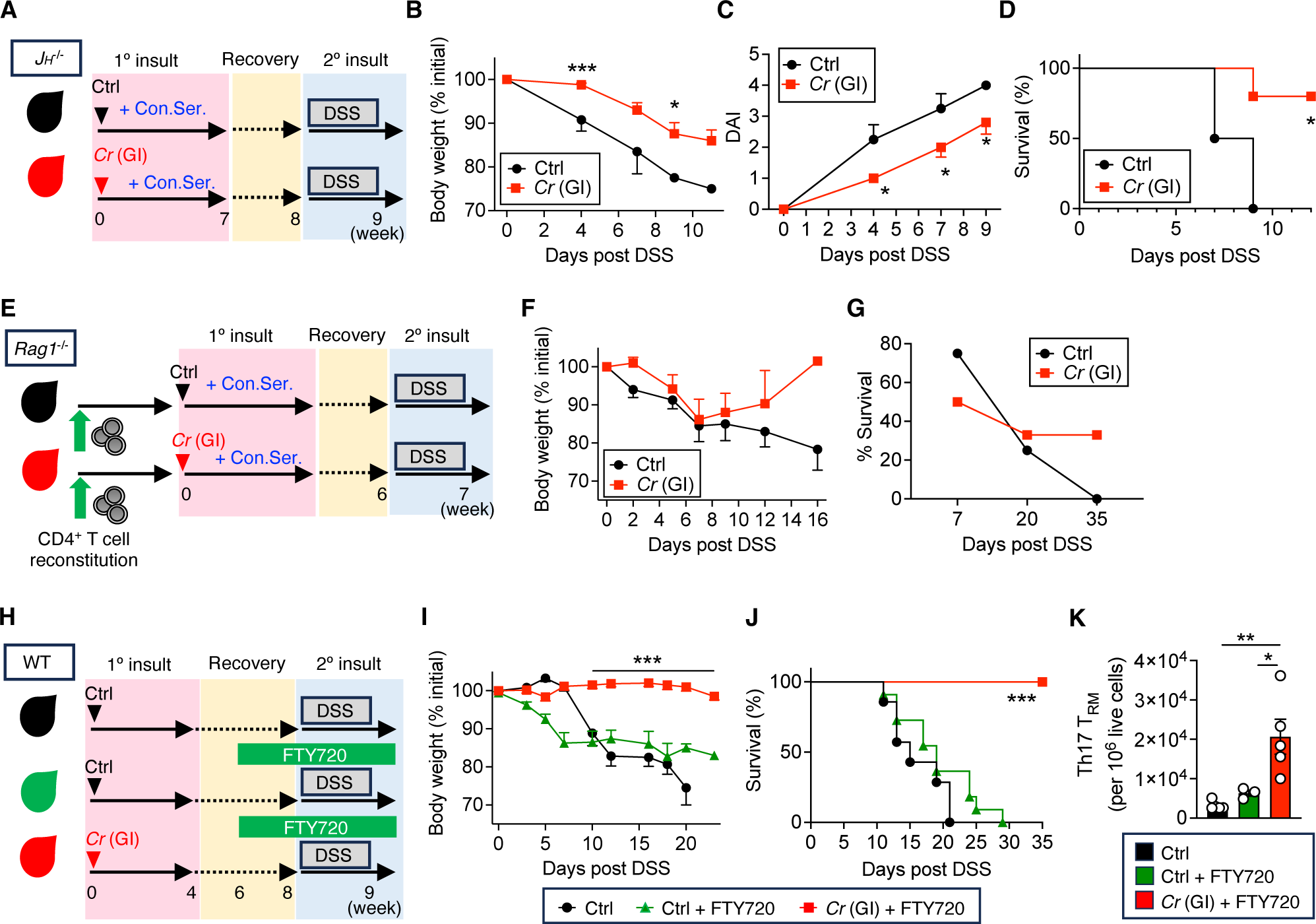
Inflammation induced T_h_17 cells are required for protection after transient inflammation. **A)** J_H_^-/-^ mice were given convalescent (con) serum from wild type mice that had cleared GI *Cr*, and subject to GI *Cr* or PBS and **(B)** weight change, **(C)** disease activity index (DAI) and (D) survival were measured. **E)** RAG1^-/-^ mice were reconstituted with wild-type total CD4^+^ T-cells and then subject to GI *Cr* or PBS. Because IgG is required to clear *Cr*, all mice were also given convalescent (con) serum from mice that had previously cleared GI *Cr*. All mice were then subject to 2.5% DSS at week 6 and **F)** weight change and **G)** survival were measured. **H)** Wild-type mice were subject to PBS or GI *Cr*, allowed to clear and then subject to FTY 720 at week 6 followed by 2.5% DSS at week 8. Control mice received PBS without FTY 720. **I)** Weight change and **(J)** survival were measured. **K)** IL-17^FM^ mice were subject to PBS or GI *Cr* +/- FTY 720 as in (H) and T_h_17 T_RM_ in the colon were enumerated at week 8 (prior to DSS). * to *** p < 0.01-0.0001 by ANOVA (B, C, F, I, K) or Kaplan-Meier (D, G, J). Data are representative of 2-6 experimental replicates, with 3-5 mice per group, per experiment.

Convalescent serum from mice that have cleared GI *Cr* contains not only anti-*Cr* IgG, but also anti-commensal IgG. We thus also considered if convalescent serum alone could mediate protection from IEC injury. To test this, we subjected WT mice to GI *Cr* or PBS and convalescent serum or non-specific control IgG at the time of DSS (**Sup Fig 3D**). In this schema, only WT mice subject to GI *Cr* have intestinal T_h_17, while the other groups would not. Consistent with our hypothesis, only mice that were subject to GI *Cr* were protected, indicating that convalescent serum alone is not protective (**Sup Fig 3E**). This result is consistent with the idea that inflammation induced T_h_17 may mediate protection.

As a second system, we used RAG1^-/-^ mice. RAG1^-/-^ mice succumb to *Cr* but can be rescued by reconstituting them with WT total CD4^+^ T-cells and administering convalescent serum from mice that have cleared *Cr* ^39^. We reconstituted RAG1^-/-^ mice, verified the reconstitution with FCS, and then subjected them to transient inflammation with GI *Cr* (**Fig 3E, Sup Fig 3F**). Control mice received PBS. All groups received convalescent serum. Mice that received CD4^+^ T-cells and convalescent serum all cleared *Cr*, verifying that reconstitution was effective. After recovery and rest, we subjected mice to injury with DSS. RAG1^-/-^ mice subjected to GI *Cr* have inflammation induced T_h_17 cells, while PBS control mice have CD4^+^ T-cells but not inflammation induced T_h_17 cells. Consistent with the possibility that inflammation induced T_h_17 are protective, higher fractions of mice that had recovered from GI *Cr* were alive at late times relative to PBS controls (**Fig 3F, G**).

Lastly, we used FTY 720, which inhibits S1PR1, to functionally isolate T_RM_. This approach is commonly employed to assess tissue-restricted T_RM_^18, 22, 47^. T-cells require S1PR1 to re-enter the circulation from tissue and lymph nodes. Circulating T-cells express S1PR1, which allows them to circulate through blood and lymph nodes^48^. In contrast, T_RM_ suppress S1PR1, which keeps them in tissues^20, 48, 49^. Blockade of S1PR1 by FTY 720 locks circulating T-cells into lymph nodes, so they cannot enter tissue, which functionally separates circulating from resident cells so that only tissue-resident cells can respond to tissue injury^25, 47, 50, 51^.

Since the protective effects of transient inflammation are tissue specific, we reasoned that if inflammation induced T_h_17 T_RM_ mediate protection, then protection would be maintained when they are functionally isolated (**Fig 3H**). As we anticipated, only mice that had recovered from transient inflammation were protected (**Fig 3I, J**), and T_h_17 T_RM_ only expanded in these mice (**Fig 3K**). Collectively, these data indicate that inflammation induced intestinal T_h_17 are required for protection.

### The inflammation-trained microbiota is required for protection

The gut microbiota is a key component of the intestinal environment that regulates immunity and exhibits memory responses ^53^. Since these features are consistent with our data, we considered the role of the microbiota in IEC protection.

After recovering from transient inflammation, GI *Cr* mice have no defects and are indistinguishable from uninfected controls. Despite returning to baseline, the gut microbiota of GI *Cr* mice is durably altered, characterized by an increased abundance of *Tenericutes* and decreased abundance of *Firmicutes*, indicating that transient inflammation leads to lasting changes in the microbiota (**Fig 4A**). To determine if this ‘inflammation-trained’ microbiota is required for IEC protection, we subjected mice to fecal microbiota transfer (FMT) with bedding transfer. We selected bedding transfer because it permits us to strictly control the direction of transfer. Mice were separated mice into four groups (**Fig 4B**). Group (1) ‘*Cr* (GI)’ which had cleared GI *Cr* and are the FMT donors. Group (2) ‘*Cr* (GI) + Abx (antibiotics)’, which had cleared GI *Cr*, then received antibiotics to disrupt the inflammation-trained microbiota. Group (3) ‘FMT recipients’, which are uninfected mice and are FMT recipients. These mice received antibiotics before FMT to increase the fidelity of FMT engraftment. Group (4) ‘Abx’, which are naive to inflammation and are controls for antibiotics. The microbiota was assessed at baseline, and week 6, 8 and 9 (**Fig 4B**).

**Figure 4.**
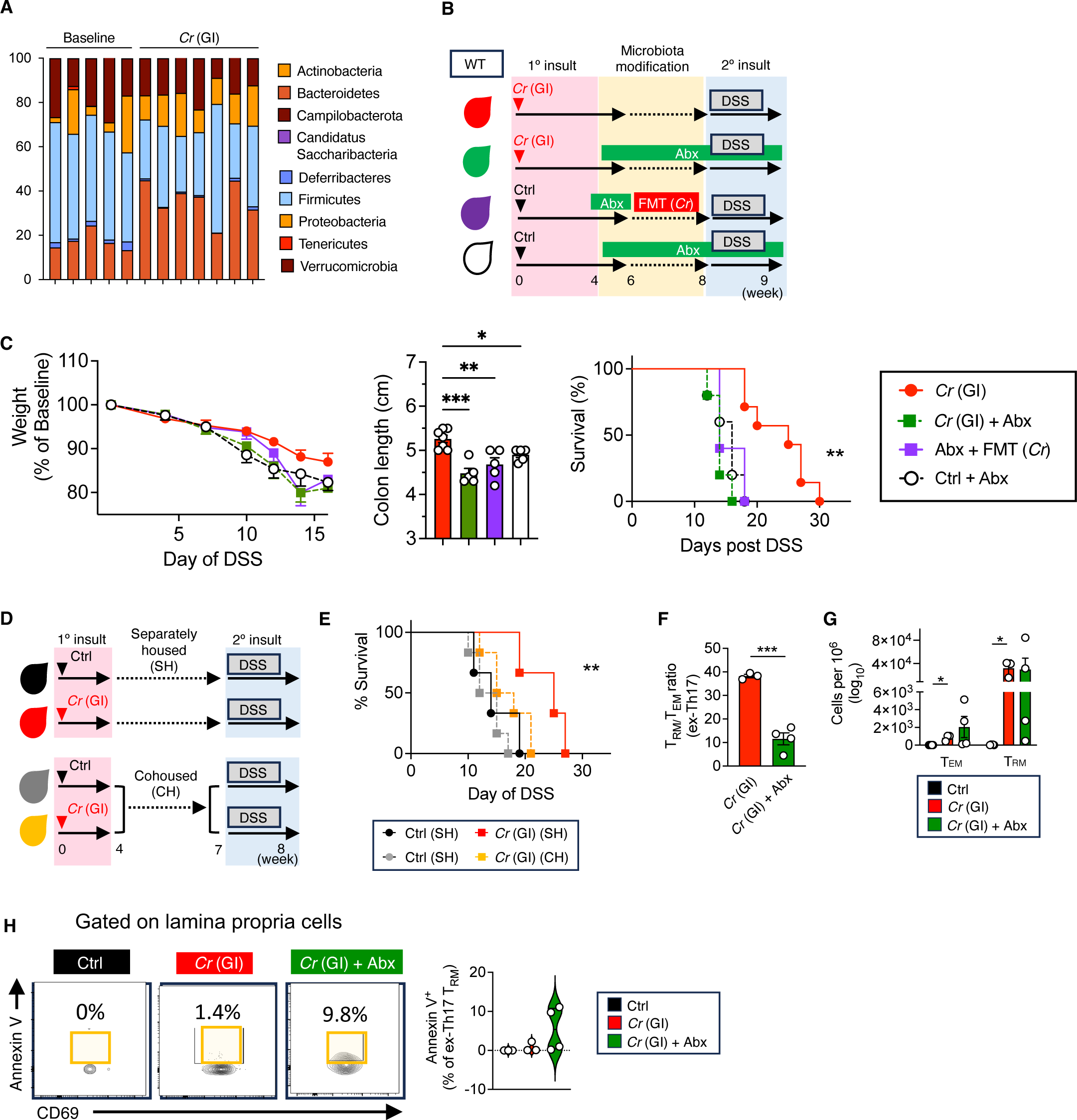
The inflammation-trained microbiota maintains inflammation induced T_h_17 T_RM_. **A)** The microbiota of IL-17^FM^ mice was assessed at baseline and 2 weeks after clearing GI *Cr*. **B)** Schematic of fecal microbiota transfer (FMT) experiments with 4 groups of mice (*Cr* (GI), *Cr* (GI) + Abx, Abx + FMT (*Cr*), and Ctrl + Abx). **C)** Weight change (left panel), colon length (middle panel) and survival (right panel) in groups from (B) with 2.5% DSS. **D)** Schematic of co-housing experiments showing separately housed (SH) and cohoused (CH) cohorts of controls (Ctrl) and *Cr* (GI) mice. After CH, mice in this cohort were separated into cages based on prior history of *Cr* or PBS. All cohorts were then subject to 2.5% DSS and **(E)** their survival was determined. **F, G)** Lamina propria mononuclear cells were assessed by flow cytometry after 2 weeks of antibiotics in the indicated groups from (B), and (F) the ratio of the fraction of CD4^+^ T_EM_/T_RM_ was quantified and (G) enumerated. **H)** Mice were left unmanipulated (Control) or subject to GI *Cr*, allowed to recover, and then monitored (GI *Cr*) or treated with 2 weeks of antibiotics (GI *Cr* + Abx). At week 8 post-infection, lamina propria mononuclear cells were assessed for expression of Annexin V by flow cytometry (left panel) and the data were quantified (right panel). The gating scheme is Live → CD3^+^CD4^+^ → GFP^+^ → CD44^+^CD69^+^ → Annexin V. * to *** p < 0.01-0.0001 by Kaplan-Meier (C, E), students t-test (F) or ANOVA (C, G). Data are representative of 3 (B, C), 2 (D-F), or one (G, H) experimental replicates with 3 to 7 mice per group.

In this protocol, only Group 1 ‘*Cr* (GI)’ have both the inflammation-trained microbiota and inflammation induced T_h_17 (**Table 1**). All other groups will lack either the inflammation-trained microbiota or inflammation induced T_h_17. If antibiotics ameliorates protection in Group 2 (‘GI *Cr* + Abx’), this means that both the inflammation-trained microbiota and inflammation induced T_h_17 are required for protection (**Fig 4B**).

**Table 1.** Schematic of the presence or absence of the inflammation-trained microbiota (left column) and inflammation induced T_h_17 (middle column) and protection from subsequent IEC injury (right column) accompanying Fig 4B.

| | Inflammation<br>trained<br>microbiota | Inflammation<br>induced<br>$T_h17$ ($T_{RM}$) | Protection |
| --- | --- | --- | --- |
| *Cr* (GI) | yes | yes | yes |
| *Cr* (GI) + Abx | no | yes | x |
| Abx + FMT (*Cr*) | yes | no | x |
| Ctrl + Abx | no | no | x |

Based on the Inverse Simpson and Shannon indices, operational taxonomic units, and family-and genus-level data, FMT from GI *Cr* successfully engrafted into recipients (**Sup Figs 4A-D**). Antibiotics effectively disrupted the SPF gut microbiota in uninfected mice but did not substantially alter the inflammation-trained microbiota in GI *Cr* mice, suggesting that that the inflammation-trained microbiota is somehow resilient (**Sup Figs 4A-D**).

Despite successful engraftment, FMT did not transfer protection (**Fig 4C**). Even more interestingly, though the inflammation-trained microbiota is resistant to major shifts with antibiotics, these mice lost protection (**Fig 4C**). We verified these results using co-housing as a second system to transfer the microbiota. As with FMT, the inflammation-trained microbiota did not transfer protection, and protection could be ameliorated by disrupting it (**Fig 4D, E**).

Given these data that the inflammation-trained microbiota and inflammation induced T_h_17 are both required for protection, we considered whether the inflammation-trained microbiota regulates inflammation induced T_h_17. Since inflammation induced T_h_17 cells exhibit a T_RM_ signature, and T_RM_ self-renew in tissue, we reasoned that the inflammation-trained microbiota may regulate the survival of T_RM_. Using FCS, we noted that there was a major shift in the ratio of T_h_17 T_RM_ to T_h_17 T_EM_, with reductions in T_h_17 T_RM_ in some mice with antibiotics (**Fig 4F, G**). Accordingly mice with reductions in inflammation induced T_h_17 T_RM_ specifically exhibited a higher fraction of Annexin V^+^ T_h_17 T_RM_, suggesting that the inflammation-trained microbiota is required to maintain these cells, at least in some circumstances (**Fig 4H**). Collectively, these data show that the both the inflammation-trained microbiota and inflammation induced CD4^+^ T_RM_ are required for protection.

## DISCUSSION

T-cells, especially T_h_17 cells, are strongly linked with IBD by murine and human studies and are considered important targets of all approved therapies for IBD ^4–8, 15, 19^. However, not all T-cells are pathogenic, and most are probably beneficial. T-regulatory (T_reg_) cell are the prototypical example of protective T-cells but there are others, including IL-10 producing non-pathogenic T_h_17 cells^33–35^. Protective T-cells may be ideal treatments for IBD because they offer the possibility of targeted therapy with little toxicity. Unfortunately, these treatment approaches have lagged, partly because the identity of protective T-cells and how they mediate protection are unclear. In exploring these questions, we inadvertently found that transient inflammation is protective from subsequent IEC injury. Our data also shows that protection is tissue specific and is encoded in the intestinal microenvironment. Based on a variety of approaches including reconstituted RAG1^-/-^ and JH^-/-^ mice, pharmacologic inhibition of cell trafficking, and FMT experiments, we found that IEC protection after transient inflammation required inflammation induced T_h_17 cells and the inflammation-trained microbiota. This finding was surprising since prior publications indicate that inflammation induced T_h_17 are pro-inflammatory ^23–25^. We thought that transient inflammation would exacerbate subsequent IEC injury due to the abundance of inflammation induced T_h_17 in the intestine.

Exactly how the inflammation-trained microbiota regulates inflammation-induced T_h_17 T_RM_ is uncertain, though our initial results suggest that the inflammation-trained microbiota maintains inflammation induced T_h_17 T_RM_. More broadly, while T_RM_ are anatomically close to the host-associated microbiota, exactly how the microbiota regulates T_RM_ is poorly understood^52^. It is unclear if the self-renewal of T_RM_ *in situ* is T_RM_ intrinsic or whether it requires microbiota derived metabolites or direct antigenic stimulation by microbes that are cross-reactive to the t-cell receptor specificity of a given T_RM_.

Tissue-resident memory T-cells were initially identified while investigating long-lived antigen specific T-cells with vaccinations and in host-defense. T_RM_ are highly enriched at muco-cutaneous body surfaces that heavily interact with microbes, like the intestine. T_RM_ can be either CD8^+^ or CD4^+^, but much less is known about CD4^+^ T_RM_ owing to fewer antigen specific murine models to track CD4^+^ T_RM_^23–25^. While T_RM_ are the leading arm of adaptive immunity *in situ*, recent data indicates that they can also be pro-inflammatory and may promote IBD. CD8^+^ T_RM_ have been linked with Ulcerative Colitis by single-cell sequencing data from intestinal specimens^31^. We found that CD4^+^ T_RM_ are enriched in Crohn’s Disease and are a major source of the pathogenic cytokine Tumor necrosis factor a^26^. Furthermore, CD4^+^ T_RM_ can drive colitis in murine models, linking them causally with the pathogenesis of IBD^32^. Thus, it is probable that T_RM_ are heterogenous and include subsets that provide host defense or are pathogenic. Adding to this, our data indicates that some T_RM_ subsets may also be protective. Our understanding of T_RM_ subsets is nascent, and one explanation for these seemingly conflicting data is that even antigen specific T_h_17 T_RM_ may be composed of cells with heterogeneous functional capacities, much like their circulating counterparts, T_h_17 T_EM_.

If some inflammation induced T_h_17 cells can be protective, why does this not appear to occur with the inflammatory injury-repair cycles of IBD? In nature, humans are constantly subject to inflammation that damages IECs. For example, fecal and food borne viral and bacterial pathogens like enterovirus and *Enterobacteraceae* are endemic and injurious to IECs. Moreover, healthy humans harbor pathobionts like *Enterobacteraceae* that are associated with IBD, but rarely develop IBD even after IEC injury and potential blooms of pathobionts. Instead, environmental insults most often generate compensatory mechanisms like adaptive immunity that make the host resilient to repetitive injury. This paradigm of inflammation leading to host resiliency makes evolutionary sense since it would confer a survival advantage to organisms. Given that IBD can manifest as cyclical injury-repair, it is possible that some IBD may be due to a breakdown of compensatory resiliency pathways.

Our data suggests a protective loop wherein transient inflammation can generate inflammation induced T_h_17 cells and inflammation-trained microbiota, which synergize to mediate protection from IEC injury. The organims has traditionally been considered as the unit upon which the environment acts. However, the fact that the host is inseperable from the host-associated microbiota has given rise to a new biological unit, the ‘meta-organism’ ^53, 54^. Thus, the meta-organism, encompasing the host and the host-associated microbiota, may be the basic unit upon which nature acts. Reframed this way, our data suggests that a primary insult can ’prime’ the meta-organism to be resilient against future injury by coupling of inflammation induced CD4^+^ T_RM_ and the inflammation-trained microbiota. In nature when insults are repetitive, this pathway fosters resiliency against subsequent injury and seems to provide a survival advantage. IBD is a cycle of repetitive injury, and one pathway for IBD may be when these compensatory systems break down. Thus, understanding these pathways will add to our scientific knowledge of intestinal homeostasis, and may lead to new therapies for IBD.

## METHODS

### Mice

Wild-type (WT) C57BL/6, RAG1^-/-^, JH^-/-^ and IL-17GFP fate-mapping (IL-17^FM^) mice were purchased from the Jackson Laboratories (Bar Harbor, ME). IL-17^FM^ mice are on a C57BL/6 background and are well published. Detailed information on the derivation of this strain can be found at the Jackson Labs website (Strain number 018472). All experiments used 5-8 wk old gender- and age-matched mice, and all mice were housed in specific-pathogen free facilities at the University of Michigan. For all experiments, mice were monitored for health by weight, spontaneous movement, diarrhea, and grooming. This protocol was approved by the University of Michigan Animal Care and Use Committee

### Citrobacter rodentium infection

Wild-type *C. rodentium* (ATCC 51459) from frozen stock was cultured in 5 ml of Luria Bertani (LB) broth supplemented with Ampicillin (100 µg/ml) overnight on a shaker at 200 rpm at 37C to generate viable colonies for infection. A spectrophotometer assessed the colony forming units (CFU)/mL after the overnight incubation. Oral inoculum stocks of 10^9^ CFU/200 µl were prepared by diluting the overnight *C. rodentium* cultures in PBS. Mice were inoculated by oral gavage with 10^9^ CFU (in 100 μl) or intravenously via tail vein injection with 5×10^7^ CFU of *C. rodentium*. Control mice received an equivalent volume of PBS via oral gavage. To quantify the burden of *C. rodentium*, the stool was weighed and then homogenized in PBS to an even suspension. Serial dilutions of the homogenized stool were plated in triplicate on MacConkey agar plates and incubated for 72 hours at 37°C. Fecal bacterial load was determined by enumerating colonies after the incubation period and was expressed per unit weight of stool (CFU/gm).

### Dextran Sodium Sulfate

2.5% DSS solution, prepared fresh, was administered to mice via drinking water for ad libidum intake as indicated until survival. Weight, grooming, consistency of stool, spontaneous movement, and rectal bleeding were monitored daily. In experiments with serial DSS, mice were administered 2.5% DSS for 7 days, and then allowed to recover. After 6 weeks, mice were subject to 2.5% DSS again and monitored for survival.

### Disease activity index, histology

Disease activity index (DAI) was scored on a scale from 0-4 based on change in body weight by increments of 5% from 0 to > 15%, qualitative assessment of stool consistency (normal to liquid with sticking to anus), and rectal bleeding (normal-gross bleeding with all stools). Histology was assessed independently by two blinded board-certified gastrointestinal pathologists (BJS, SB), and was scored from 0-4 based on epithelial damage, neutrophil predominant acute inflammation, and chronic inflammatory changes (B- and plasma-cell clusters). For both DAI and Histology, higher scores indicate worse damage. Quantitative PCR was performed on colon tissue using kit instructions.

### Quantitative PCR

Colon was homogenized in RLT buffer (RNAeasy kit, Qiagen, Valencia, CA) with a tissue homogenizer. RNA was extracted with RNAeasy kits and cDNA synthesized with iScript cDNA Synthesis kits (Hercules, California). Relative quantification of genes was determined by real-time PCR with SYBR Green normalized to GAPDH. Some data as additionally normalized to baseline conditions as noted in the Figure legends. All SYBR Green and primers were from Bio-Rad (Hercules, CA). Results were analyzed on a CFX Connect system (Bio-Rad).

### Isolation of murine and intestinal and circulating T-cells

Mouse lamina propria mononuclear cells (LPMCs) were isolated with minor modifications to this published protocol. Briefly, the colon was cleared of stool by flushing with cold Hank’s balanced salt solution (HBSS), then washed again with fresh cold HBSS. The tissue was then transferred to a ‘pre-digestion buffer’ composed of warmed HBSS, 2.5% fetal bovine serum (FBS), 5mM EDTA, and 1 mM dithiothreitol, and dissociated using the gentleMACS Dissociator (1 min, Tissue dissociating setting; Mintenyi Biotech, Bregisch Gladbach, Germany). The tissue was then incubated for 20 min at 37C with gentle mixing and was then vortexed (3,000 RPMs, 10s) and washed in fresh cold RMPI-1640. The tissue was transferred to 50 mL conical tubes with 5 mL of digestion solution pre-warmed to 37C and composed of 150 U/ml Collagenase type III (Worthington Biochemical, Freehold, NJ) and 50 µg/ml DNase I, 2% FBS in complete RPMI-1640, and incubated at 37C in a shaker for 30 min (human tissue) or 45 min (mouse tissue). The digested tissue was put through a 100 μm cell strainer, washed in cold HBSS, and centrifuged (450g, 10min, 4C), and the pellet was resuspended in 1 mL of 40% Percoll, and added to a fresh conical with 4 mL of 60% Percoll. The solution was underlayed with 2.5 mL of 80% Percoll to create a 40/80 interface. The samples were centrifuged (860g, 20 min, 21C) and the LPMCs were aspirated from the 40/80 interface. The LPMCs were washed twice with HBSS and resuspended in 1640 RPMI.

### Flow cytometry

Cells were rested overnight after purification or stimulation per above in RPMI supplemented with glutamine, sodium pyruvate, 100 units/ml penicillin, 100 µg/ml streptomycin, and 10% fetal bovine serum before stimulation with 1X stimulation cocktail (eBiosciences, Waltham MA) with Golgi Plug (BD Biosciences, Franklin Lakes, NJ) before flow cytometry for 4-6 hours. Flow cytometry was then performed on the LSR II (BD Biosciences, Franklin Lakes, NJ) or the FACS ARIA II (BD Biosciences), and the data were analyzed with FlowJo (Ashland, OR). The following antibodies were used for flow cytometry: CD3ε BV 510 (BioLegend, clone 145-2C11), CD44 BV 421 (BioLegend, clone IM7), CD8β PerCP-eFluor 710 (eBiosciences, clone H35-17.2), CD8β PerCP Cy5.5 (BioLegend, clone YTS156.7.7), IFNγ APC (BioLegend, clone XMG1.2), IFNγ PE (BioLegend, clone XMG1.2), CD4 FITC (Thermo Fisher, Waltham, MA, Clone GK1.5), CD4 PE Dazzle (BioLegend, clone GK1.5), CD4 BV 650 (BioLegend, clone RM4-5), CD69 PE-Cy7 (BioLegend, clone H1.2F3), IL-17A PE, IL-17A APC, IL-17 FITC (BioLegend, clone TC11-18H10.1), CD45 BV 605 (BioLegend, clone 30-F11), Annexin V CF405M (Biorbyt), Fix Viability eFluor 780 (Thermo Fisher)

### Reconstitution of mice with CD4^+^ T-cells

Total CD4^+^ T-cells were obtained by negative selection from the spleen of wild-type mice using EasySep^TM^ magnetic separation per kit instructions (product number 19851, STEMCELL Technologies, Vancouver, Canada). Purified total CD4^+^ were enumerated and 5×10^5^ total CD4^+^ T-cells were injected via tail vein to recipient mice. CD4^+^ T-cell purity and engraftment were verified using flow cytometry of recipient mice.

### Administration of convalescent serum

Wild-type mice were subject to GI *C. rodentium*, and 3-4 weeks post-infection, convalescent serum was collected from these mice. Recipient mice received 75-100 μl of convalescent serum (per mouse), by intraperitoneal injection, on days +3, +4 and +5 of *C. rodentium* infection. The concentration of circulating convalescent serum in recipient mice was determined with optical densitometry.

### Administration of FTY 720

FTY 720 (Cayman Chemical, Ann Arbor, MI) was mixed with drinking water to deliver an approximate dose of 3 mg/kg (body weight)/day ad libidum for 2 weeks, before 2.5% DSS or prior to sacrifice for mucosal T-cell assessments. The quantity of FTY 720 dissolved in drinking water was estimated using the average weight of mice in each cage and the total volume of the drinking water.

### Fecal microbiota transfer (FMT) and co-housing experiments

FMT experiments were done via bedding transfer. Groups of mice as indicated in the figure legends (**Fig 4B**) were kept in sperate cages without mixing throughout the FMT experiment. All groups except the bedding donor cage received 1 mg/gm (body weight) of Streptomycin (RPI, Mount Prospect, IL) via oral gavage prior to bedding transfer to facilitate FMT engraftment or as controls. FMT recipients and control mice received 2 doses of streptomycin on day -2 and -1 before bedding transfer on day 0. FMT recipients and control mice received no further antibiotics. The group of mice that had previously cleared GI *C. rodentium* received antibiotics every 3 days for the duration of the experiment, as noted in the figure legends.

For co-housing experiments, 2 cages of mice were allowed to clear GI *C. rodentium* and return to baseline. 2 cages of control mice were treated with PBS. Six weeks p.i., after mice had returned to baseline, mice that had recovered from GI *C. rodentium* were co-housed with control PBS gavaged mice for 3 weeks. To minimize cage effects, mice were chosen at random from each group (GI *C. rodentium* or PBS) for co-housing. Mice from cages that were not selected for co-housing were treated as non-cohoused controls, as indicated in the figure legends. After 3 weeks, co-housed mice were separated into cages based on whether they had cleared GI *C. rodentium* or were PBS gavaged controls, and all cages were subject to 2.5% DSS.

### Other Statistics

Data were analyzed using the ANOVA, unpaired students *t*-tests or with Mann-Whitney correction as noted in the Figure legends. All data were analyzed using Prism (GraphPad, San Diego, USA).

### Quantification of 16s for microbial taxonomy

For 16s gut microbiome analysis, samples were collected from each mouse at four different time points as indicated in Fig 4B and stored at -80C before further processing. Fecal genomic DNA was extracted with the DNeasy Blood and Tissue Kit (QIAGEN, Germantown, MD) using a previously published protocol that was modified slightly for optimization ^55^. The extracted DNA was quantified using Nanodrop. Specimens the underwent PCR amplification of the V4 region of the 16S rRNA gene small subunit microbial gene (‘16S’) with the EMP primers ^56^, and the libraries were sequenced on the Illumina MiSeq platform as per the standard protocol of the University of Michigan Microbiome Core.

The resultant raw reads were then processed with MaLiAmPi ^57^, a Nextflow-based ^58^ pipeline for the pre-processing and phylogenetic placement of 16S data. In brief, primers and adapters were removed with TrimGalore and then reads had error deconvolution via DADA2 into amplicon sequence variants (ASVs) after filtering and trimming ^59^. The ASVs were then phylogenetically placed with pplacer on a tailored phylogenetic tree of full-length 16S rRNA SSU gene alleles recruited from the YA16SDB repository of alleles ^60^. The guppy package (a part of the pplacer suite) was then used to estimate alpha-diversity for each specimen and assign taxonomy to each ASV based on the phylogenetic placement generated. These tables were then used for subsequent analysis.

### Statistical analysis of microbial taxonomy

For Fig 4A, stacked bar plots were generated based on specimen ASV counts normalized to relative abundance and then ordered and colored by the taxonomic assignments from the hybrid2 classifier in guppy. Operational taxonomic unit assignments and diversity measurements for Sup Fig 4A-D were performed using mothur via the microbiome core of the University of Michigan as previously published ^61^. All analysis was performed using R.

## Acknowledgments

Min Zhang for supporting experiments, and the University of Michigan Microbiome Core for data analysis.

---

**Supplemental Figure 1.**
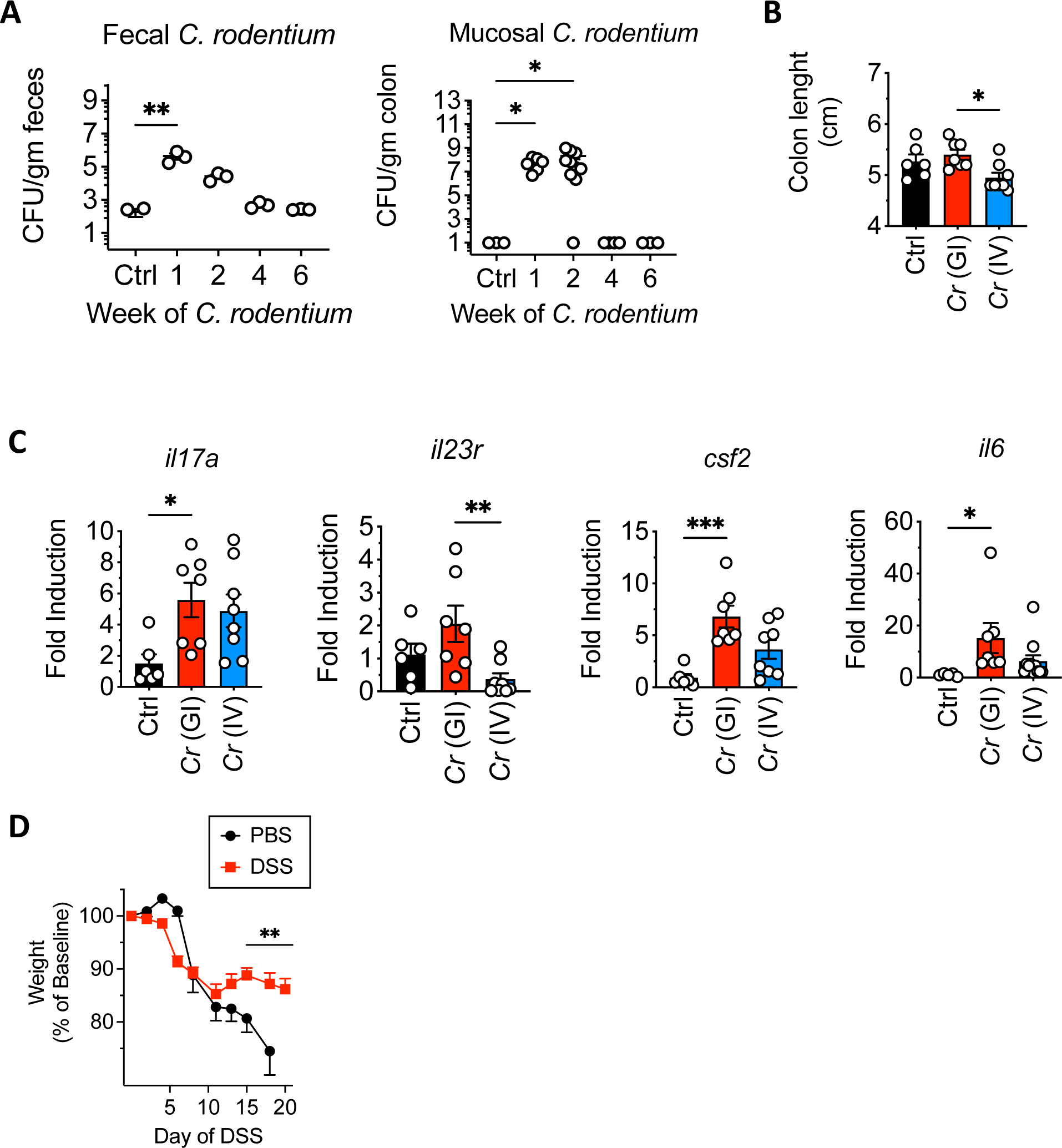
**A)** Mice were subject to GI *Cr* and colony forming units (CFU) from feces and the colon (left and right panels, respectively) were enumerated at the indicated times. Control (Ctrl) are mice at baseline. Mice were subject to 2.5% DSS after recovering from GI or IV *Cr* or PBS as in Fig 1A and **(B)** colon length and **(C)** gene transcripts from the colon were measured at day 10 of DSS. **D)** Weight loss of wild-type mice subject to sequential DSS or PBS as in Fig 1F. * to *** p < 0.01-0.0001 by ANOVA. Data are representative (A, D) or cumulative (B, C) of 3-6 experimental replicates, with 3-5 mice per group, per experiment.

**Supplemental Figure 2.**
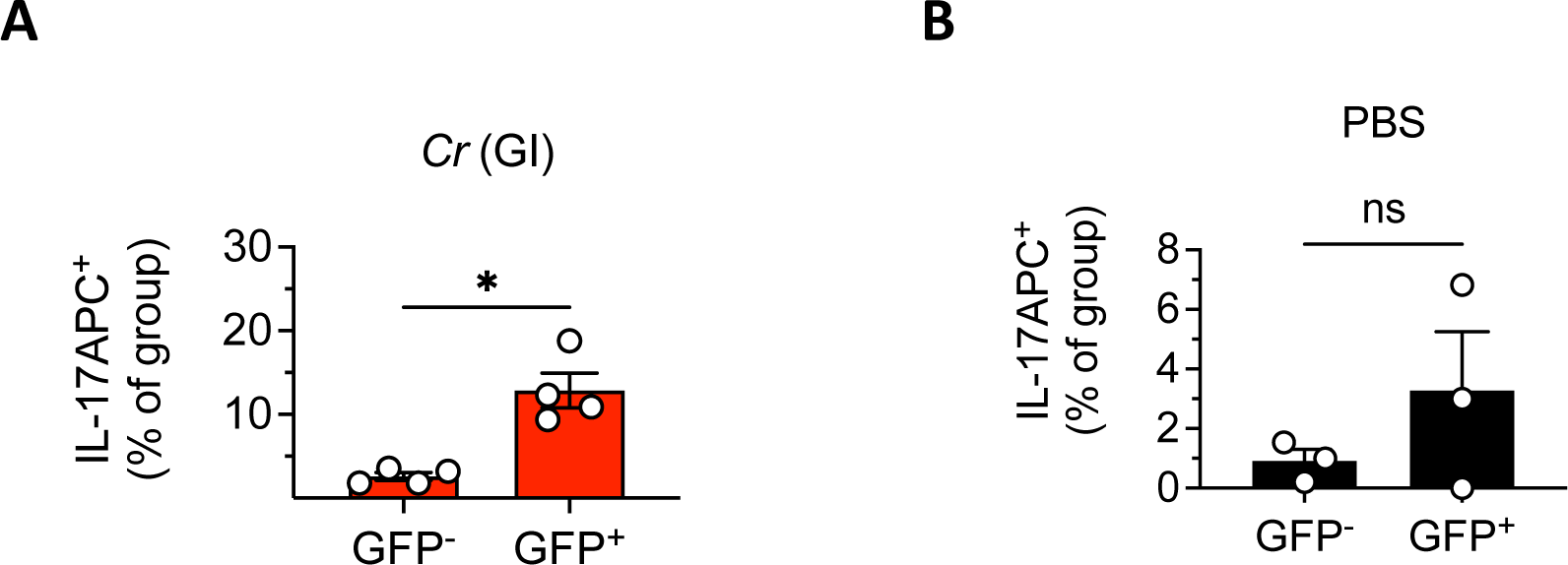
Lamina propria mononuclear cells from the colon of IL-17^FM^ mice 2 weeks after recovering from **(A)** GI *Cr* or **(B)** in PBS controls as in Fig 2A were counter-stained with IL-17APC. Lamina propria cells were then gated on CD3^+^CD4^+^GFP^+^ (T_h_17) and CD3^+^CD4^+^GFP^-^ (non-T_h_17) and the fraction of IL-17APC^+^ cells in these groups was quantified. * p < 0.01 by students t-test. Data are representative of 2 experimental replicates, with 3-4 mice per group, per experiment.

**Supplemental Figure 3.**
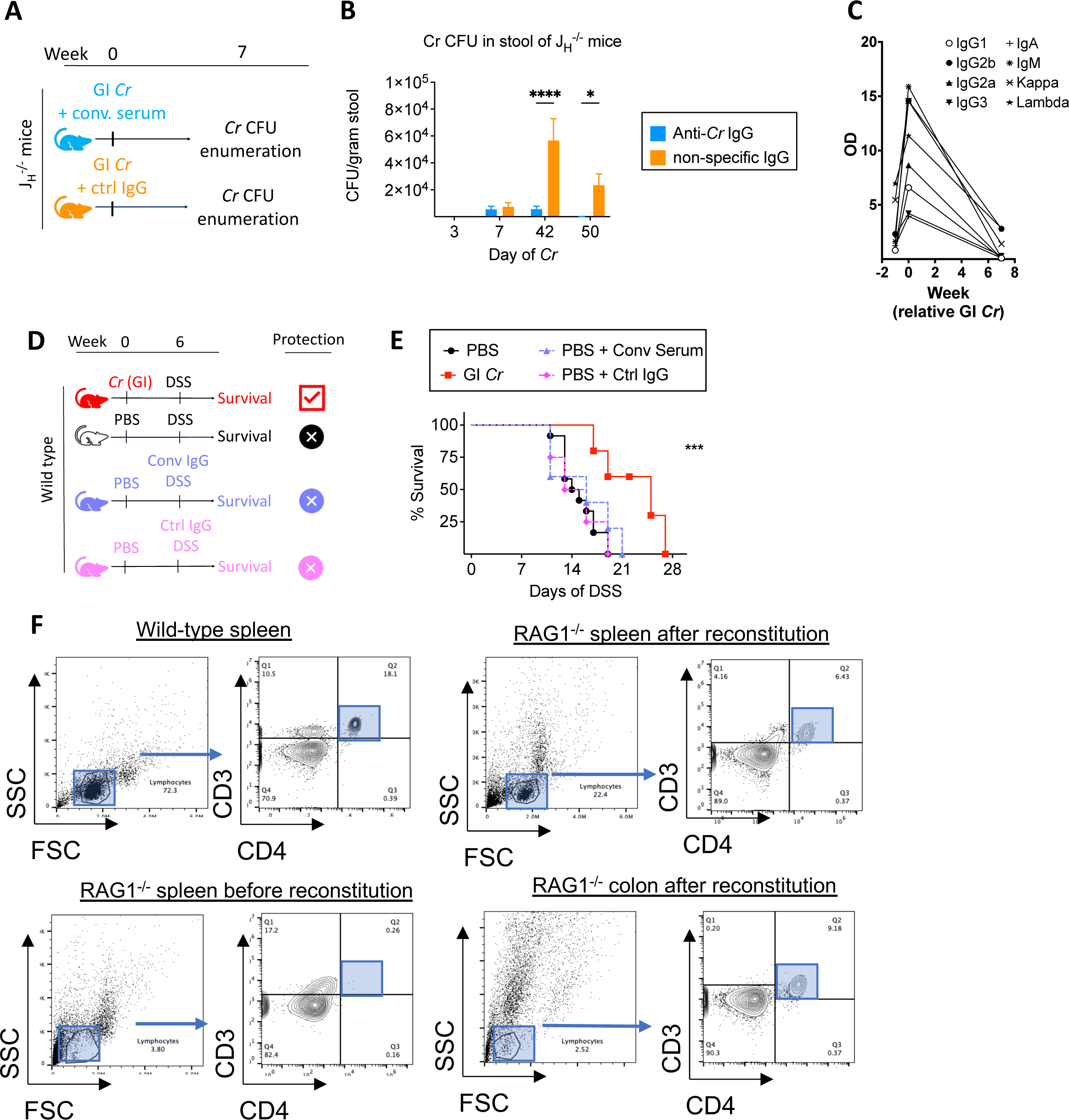
**A)** J_H_^-/-^ mice were given convalescent (conv) serum from wild-type that had cleared GI *Cr* or control (Ctrl) IgG and both groups were subject to GI *Cr* and colony forming units (CFUs) of *Cr* from feces was **(B)** enumerated at the indicated times. **C)** The concentration of convalescent serum was quantitated from the tail vein by optical density (OD) at the indicated times relative to GI *Cr* infection. **D)** Wild-type mice were subject to GI *Cr* or PBS followed by 2.5% DSS with convalescent (conv) serum or control (Ctrl) IgG as depicted and **E)** survival was measured. **F)** Flow cytometry analysis of CD4^+^ T-cells in the spleen of (top left panel) wild-type and (bottom left panel) RAG1^-/-^ mice before reconstitution with total CD4^+^ T-cells and in the spleen and colon of RAG1^-/-^ mice (right top and bottom panels, respectively) after reconstitution with total CD4^+^ T-cells. * to *** p < 0.01-0.0001 by students t-test (B) or Kaplan-Meier (E). Data are representative of 1-2 experiments (A-C, F) or cumulative from 2 experiments (D, E) with 3-8 mice per group.

**Supplemental Figure 4.**
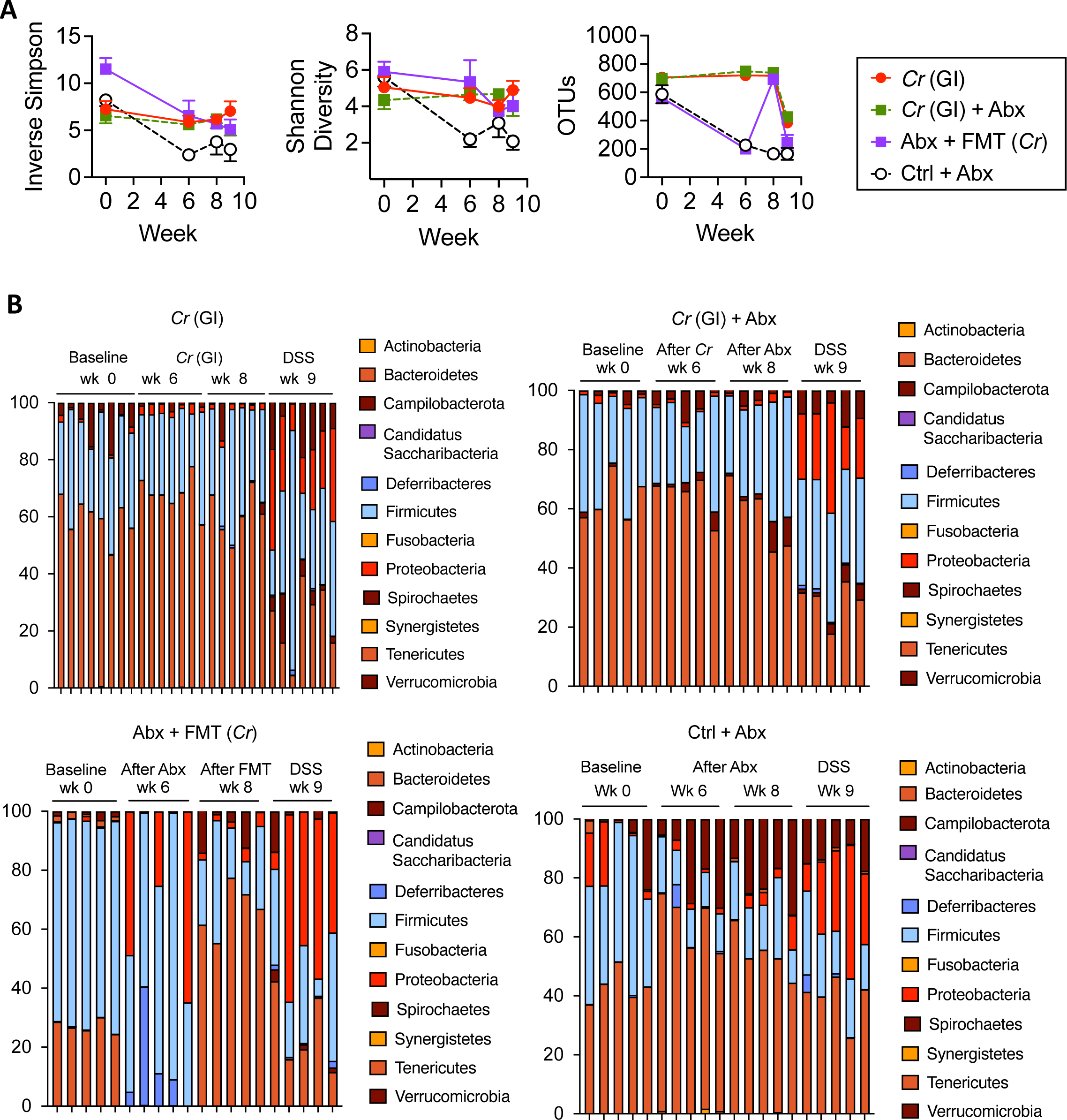
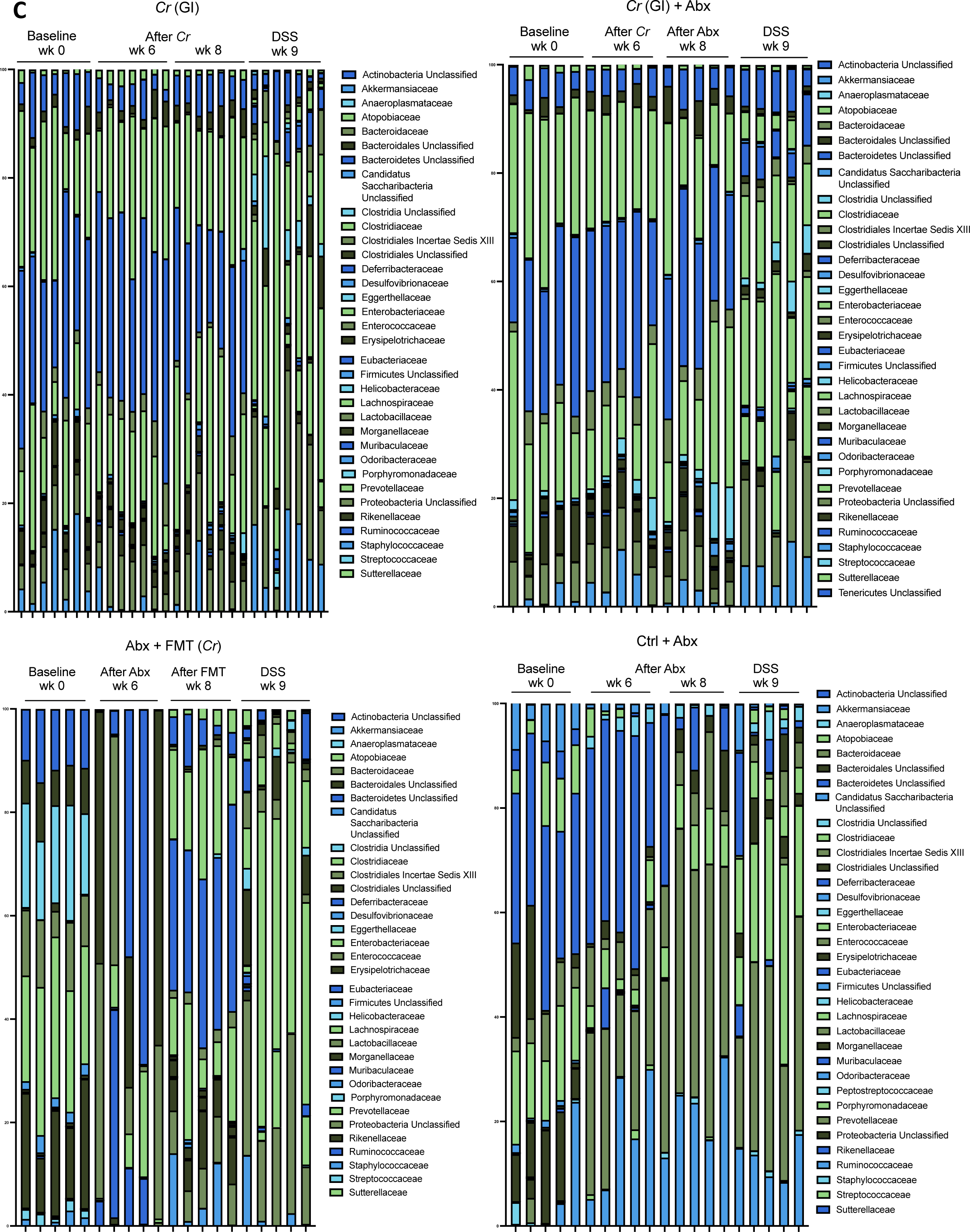
**A)** The Inverse Simpson (left panel) and Shannon (middle panel) indices and operational taxonomic units (OTUs) (right panel) from mice in Fig 4B. **B)** Phylum and **(C)** family level data from mice in Fig 4B. Data are representative of 3 experiments, with 3-5 mice in each group, per experiment (12-20 mice per experimental replicate).

